# Rapid flapping and fiber-reinforced membrane wings are key to high-performance bat flight

**DOI:** 10.1101/2023.09.11.557136

**Authors:** Marin Lauber, Gabriel D. Weymouth, Georges Limbert

## Abstract

Bats fly using significantly different wing motions than other fliers, stemming from the complex interplay of their membrane wings’ motion and structural properties. Biological studies show that many bats fly at Strouhal numbers, the ratio of flapping to flight speed, 50-150% above the range typically associated with optimal locomotion. We use high-resolution fluid-structure interaction simulations of a bat wing to independently study the role of kinematics and material/structural properties on aerodynamic performance and show that peak propulsive and lift efficiencies for a bat-like wing motion require flapping 66% faster than for a symmetric motion, agreeing with the increased flapping frequency observed in zoological studies. In addition, we find that reduced membrane stiffness is associated with improved propulsive efficiency until the membrane flutters, but that incorporating microstructural anisotropy arising from biological fiber reinforcement enables a tenfold reduction of the flutter energy whilst maintaining high aerodynamic efficiency. Our results indicate that animals with specialized flapping motions may have correspondingly specialized flapping speeds, in contrast to arguments for a universally efficient Strouhal range. Additionally, our study demonstrates the significant role that the microstructural constitutive properties of the membrane wing of a bat can have on its propulsive performance.

---

Bats are amazing fliers able to perform powered flight using membrane wings which endow them with exceptional maneuvering capabilities [1]. Understanding the complex mechanical interplay of their membrane wing and flight kinematics is key to unraveling fundamental questions in evolutionary biology and to engineering of biologically inspired flying vehicles.

Numerous studies have investigated the kinematics of bats in forward flight [2–7] and provided details of its unique features. Bats fly using a power stoke where the aerodynamic forces are mainly produced during the downstroke while the upstroke is comparatively far less active and can even be feathered [8, 9]. The downstroke starts with the wing moving ventrally and anteriorly along the stroke plane (see Fig. 1A). An essential parameter of bat (and animal) flight is the angle of the stroke plane relative to the horizontal, which tends to increase with flight speed [1]. Wing extension is also a key influencing parameter of bat flight; it is maximal during the downstroke to enhance aerodynamic force production [10] while it is minimal during the upstroke to reduce power expenditure [11]. An aspect that differentiates bat flight from other flying animals is their wing’s weight, representing up to 20% of their total weight [12], which influences the power requirements of bats. While these studies provide excellent observation of bat flight, they cannot isolate the essential aspects of kinematics and their influence on flight efficiency. Additionally, reported flapping frequencies across numerous bat species can vary by as much as 50% compared to those of morphologically similar birds, at equivalent flight speed and flapping amplitude [13, 15]. We compile the experimental measurements of [13] (see SI Appendix Tab. S1) in terms of the Strouhal number *St*

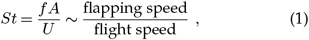

with *f* the flapping frequency, *U* the forward velocity of the swimmer/flyer, and *A* is the peak-to-peak flapping amplitude, see Fig. 1A. Animal flight or swimming typically occurs in a narrow range of Strouhal numbers, 0.25 *< St <* 0.35, and this narrow band is linked to optimal locomotion in terms of propulsive efficiency [16] associated with the most unstable-mode of the wake. However, Fig. 1B shows that many bats fly at Strouhal numbers significantly higher than this, and we find that the motion amplitude is a good measure of wake width, see SI Appendix Fig. S1, meaning our use of *A* in defining *St* is consistent. As such, the anomaly in the Strouhal number indicates that bat flight is an outlier compared to other flapping animals.

**Figure 1.**
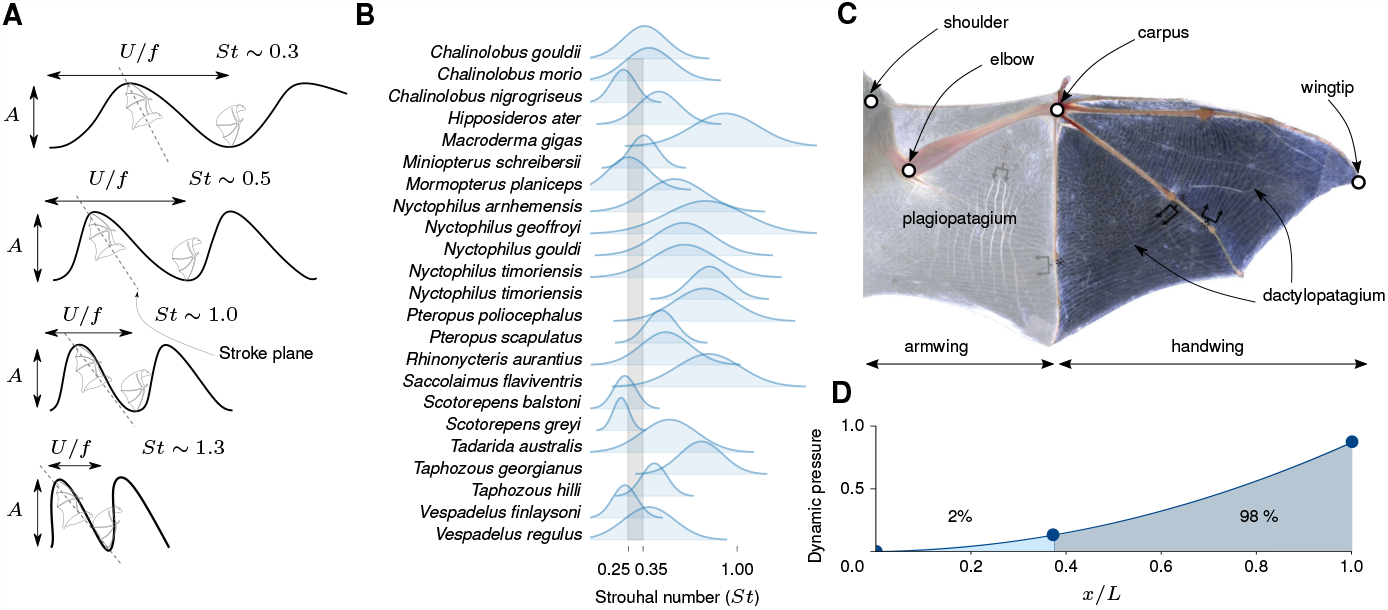
Schematic of **A** the Strouhal number of a bat with showing the wingtip trajectory (black line) and the stroke plane (dashed grey line), adapted from [1]. **B** Strouhal number envelope for several bat species from the data provided in [13]. The normal distribution is drawn with a variance where the min-max envelope corresponds to *±*3*σ* (99.7% confidence interval), and the mean uses the mean Strouhal number from the data. The grey shaded area represents the typical Strouhal range of 0.25 *< St <* 0.35. **C** Macroscopic structure of a bat wing membrane. Asterisks indicate the array of nearly homogeneous, approximately spanwise-oriented *elastin* fibers; dagger indicates the chordwise-oriented *musclesin* in the armwing, adapted from [14]. **D** Distribution of relative pressure due to dynamic motion along the wing’s span computed from experimental measurements[4].

In addition to bat flight’s uniquely high Strouhal number, bat wing material properties are also unique, arising from a complex arrangement of various types of elastic fibers and muscles. Typical membrane thicknesses range between 130 and 300 *μ*m, making them extremely thin compared to other mammals’ skin [17]. Fiber assemblies in the *plagiopatagium* and the *dactylopatagium* form an orthogonal net [17], with the fibers approaching the digits at 90^*°*^ to the digit’ long axis. These fiber arrangements are volumetrically dominated by spanwise *elastin* fibers featuring a high degree of elastic recoil, while the much stiffer *collagen* fibers are present in much smaller proportions. Within this orthogonal fiber network, there is a significant stiffness ratio between the spanwise and chordwise mechanical properties [18], see Fig. 2C. The high level of mechanical anisotropy of the wing results from pre-stretched spanwise elastin fibers embedded in a matrix with randomly oriented collagen fibers [14]. At rest, the pre-stretched elastin fibers induce buckling of the supporting matrix. Under tension, the much stiffer matrix and collagen fibers increase spanwise stiffness once the elastin fibers have been stretched past the unwrinkled configuration of the membrane. While it has been hypothesized that this fibrous net improves flight efficiency [19, 20], this has not been verified nor quantified with parametric studies rooted in continuum mechanics of both fluid and solid media.

**Figure 2.**
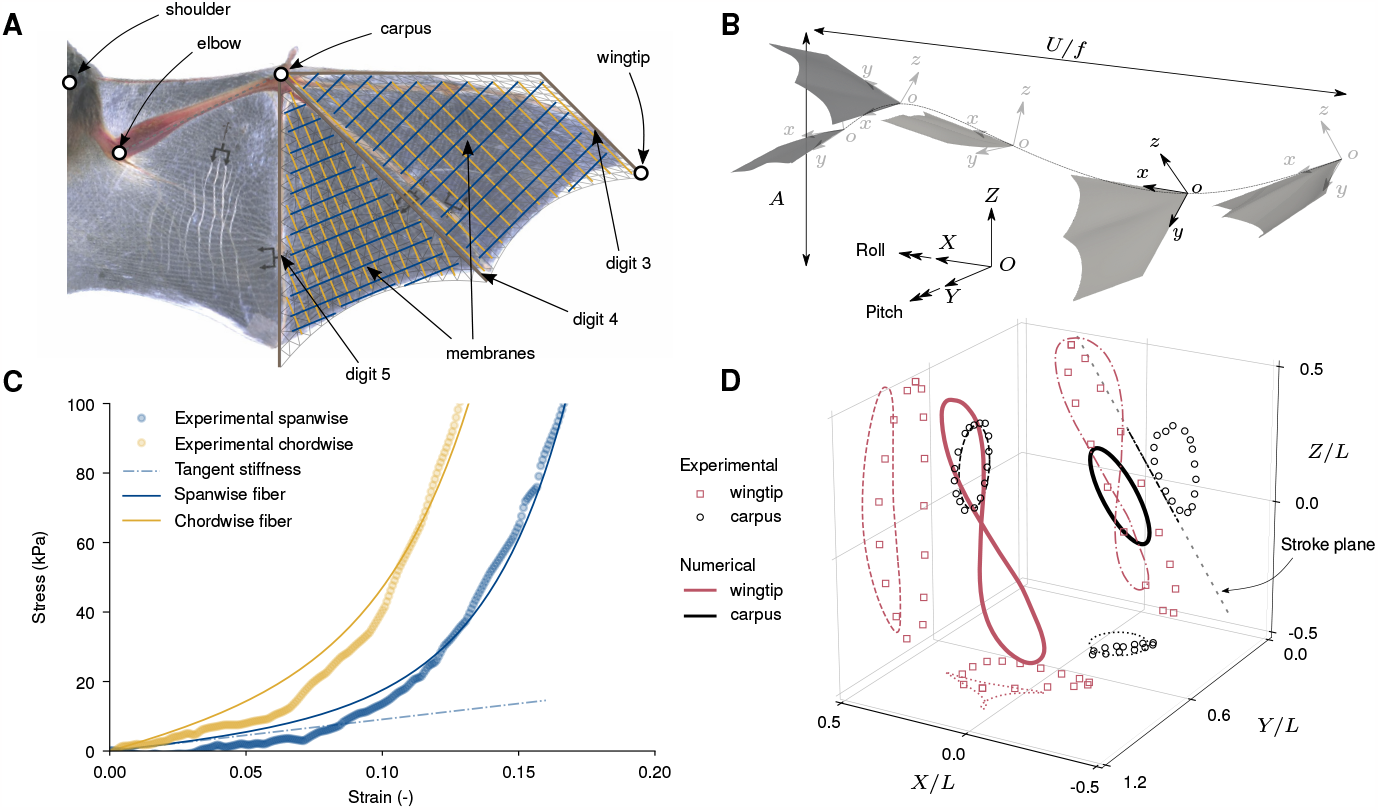
Schematic of the kinematic and geometrical model of the bat wing. **A** Coarse representation of the finite-element model of the handwing used for the numerical simulations with the fiber family direction vector. **B** Cartesian (*OXY Z*) and carpus (*oxyz*) coordinate system. The carpus coordinate system is convected with the wing following the dashed path. Pitch and roll are rotations around the *Y* and *X*-axis, respectively. The Strouhal number, *St* = *A/*(*U/f*) is depicted for kinematics at *St* = 0.5. **C** Constitutive curves for the fiber-reinforced membrane under biaxial loading showing the membrane’s non-linear behavior and the tangent stiffness modulus for calibrating the mechanical properties. The constitutive curves correspond to the two fiber families in A. The tangent stiffness modulus shows a response equivalent to linear isotropic material. **D** Trajectory of the wingtip and carpus in a 3D space (thick lines) with projections onto the three planes (various discontinuous lines) for a Strouhal number *St* = 0.5 compared to experimental measurements (markers) of [4]. The stroke plane is represented by the dashed grey line. The experimental Strouhal number is estimated to be *St ∼* 0.5.

Experimental studies of bat flight are limited in the amount of information that can be simultaneously collected about the structural and material properties of the wing, and motion of the bat during flight. Therefore, these studies cannot segregate the respective role of particular structural elements of the membrane/wing composition or that of flight kinematics on the bat’s performance. Similarly, previous numerical simulations rely either on marker measurements, *fixed* and *discrete* by definition, and one-way fluid-structure interaction coupling [10, 21], or on overly simplistic kinematics and constitutive laws for membrane skin [22, 23]. In the first case, these numerical studies inherit the experimental study’s *fixed* set of parameters by design, thus preventing any further wider parametric studies, and in the second case, the respective influence of the microstructural constitutive parameters and motions of the wing membrane, and their interplay, are not captured because of the low fidelity of the kinematics and material law used. The research presented here uses fully-coupled fluid-structure simulations to perform the first controlled study of bat performance using realistic parametric models of the bat flight motion in concert with state-of-the-art microstructurally-based constitutive model of the wing membrane valid for arbitrary (i.e. *finite*) deformation. Computational studies are indispensable for these complex systems that are intractable analytically [24]. By investigating the flapping Strouhal number and membrane reinforcement independently, we determine their direct influence on the flow field and wing deformations and ultimately explain the influence of these unique features on bat flight performance.

## Results

### Parametric modelling approach

Our geometrical model of the bat wing consists of the three distal digits (3, 4 and 5) and the membranes that make up the handwing, see Fig. 2A. The handwing experiences 98% of the dynamic load during a cycle (see Fig. 1D) and significantly contributes to both lift and thrust. Previous quasi-steady models of bat flight [12] estimate that thrust is produced mostly by the handwing while lift contributions are shared equally between the hand and armwing. However, the armwing relies on complex muscle activity to generate lift by modulating the camber of the (*plagiopatagium*) membrane [20], while the handwing relies on the large apparent velocity induced by the motion and camber modulation via folding of the digits. Attempting to actively control the time-dependent muscle and digit activity would add significant complexity and uncertainty to our geometric and kinematic model of the wing. While omitting part of the biological system such as the body and secondary lifting surfaces certainly introduces errors, focusing on the dominant contributor to the propulsive force has been used in many successful numerical studies [25, 26]. We therefore, limit the current study to the handwing with inactivated digits and focus on the dynamic thrust and lift production and passive deformation of that structure. The flight kinematics are imposed on our geometrical model through the *carpus*, see Fig.2A.

Bats use complex pitching, folding, and active camber control of their wing during the cycle to modulate the thrust/lift generation. Lift is mainly produced in the downstroke, and the upstroke has a minor contribution to the cycle-averaged values, while thrust is produced throughout the cycle [10]. Mimicking the loading applied during the downstroke and upstroke is critical to achieving realistic performance estimates (see SI Appendix Fig. S2), but modeling the folding of the wing during the upstroke is extremely complex. To provide simplified yet realistic kinematics, we prescribe a variable pitch to our model based on an effective angle of attack approach [27, 28]. This is an approximate method used to quickly estimate appropriate pitch kinematics, after which the full fluid-structure interaction solver is used to determine the resulting flight performance. However, this simple approach captures bat wing motion accurately when compared to actual wing motion [4], see Fig. 2D. The complete kinematics, their calibration, and the procedure used to obtain the pitch profiles are presented in the Method section.

We use our fully-coupled fluid-structure interaction solver, presented in [29, 30], to perform numerical simulations of our parametric geometric and kinematic model of a bat wing, see Fig. 2A and the Method section for more details. In addition to Strouhal *St*, three non-dimensional parameters govern the coupled problem: the Reynolds number, *Re* = *Ub/ν*_*a*_, the mass ratio, *M*_*ρ*_ = *ρ*_*s*_*h/ρ*_*a*_*b*, and the Cauchy number,

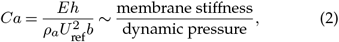

where *ν*_*a*_, *b, E*, and *h* are the kinematic viscosity of air, the bat’s half-span, Young’s modulus, and the membrane thickness, respectively. *ρ*_*s*_ and *ρ*_*a*_ are the membrane and air density, respectively. The velocity scale, *U*_ref_, is the maximum apparent velocity during the cycle 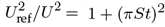. This *Ca* scaling ensures that the impact of Strouhal on the relative stiffness of the wing is accounted for, allowing us to study the relative speed and stiffness effects independently. Using typical bat wing membrane density [17], the mass ratio is set to *M*_*ρ*_ = 0.589. The full viscous flow equations are solved, and assuming a standard cruising speed and a wingspan of bats, the Reynolds number is set to *Re* = 10^4^. We study the Strouhal range *St* = 0.3 *−* 0.7 based on Fig 1B. A simple estimate of bat Cauchy number assuming a membrane wing with a thickness of 150 *μm* [17] and a wing span of 0.1 *m*, a reference (max apparent) velocity of 10 *m/s* and a tangent stiffness modulus 0.250 G*Pa* [18] gives *Ca ∼* 0.3, and we simulate a wide range of *Ca* around this estimate.

We start by presenting results for a linear isotropic elastic constitutive law to investigate the effect of the flapping speed and stiffness of the membrane wing on the aerodynamic efficiencies. We will then compare the isotropic linear elastic model to the non-linear fiber-reinforced formulation that incorporates microstructural and loading-induced anisotropy, more closely resembling actual characteristics of the bat wing’s membrane [18, 31]. Idealized responses of this fiber-reinforced model are shown in Fig. 2C together with the experimental calibration data [18]. The strain energy density function of the non-linear fiber-reinforced formulation and the method used to calibrate the material coefficients on the experimental bat membrane data are presented in the Method section.

### Flow field characterization

Figure 3 shows the vortex structures generated by the wing at three different Strouhal numbers and three different Cauchy numbers at the same time during the downstroke (*t/T* = 0.78, with *T* the motion period). These visualizations show that the wake is sensitive to both the Strouhal number and the Cauchy number of the membrane. The complete unsteady flow field evolution can be viewed in the electronic supplementary material video.

**Figure 3.**
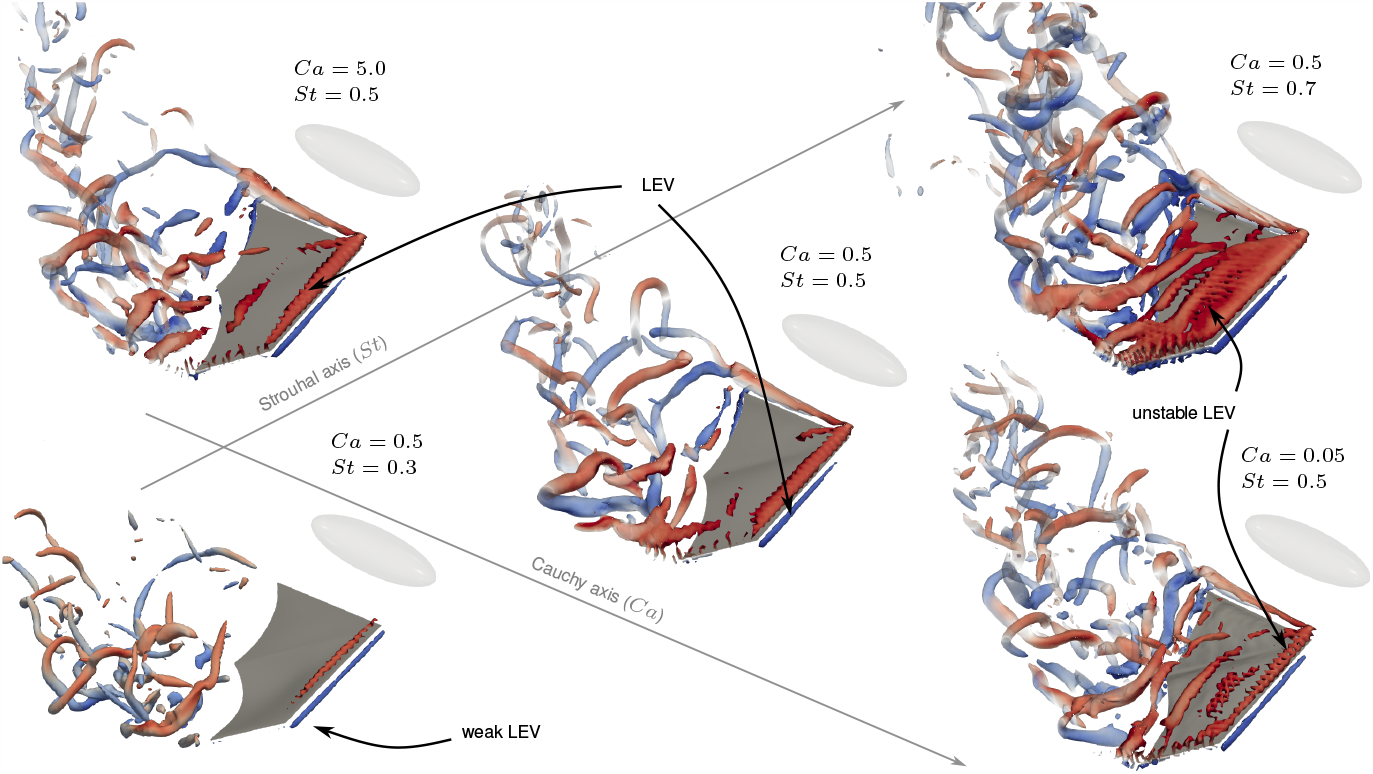
Vortex structures in the wake of the bat represented by isosurfaces of *λ*_2_(*b/U*)^2^ = *−*5 *×* 10^*−*6^ coloured by spanwise vorticity *ω*_y_(*b/U*) *±* 50 at a selected time during the downstroke. Three different Strouhal numbers at a fixed Cauchy number *Ca* = 0.5 are represented on the bottom-left to top-right axis and three different membrane Cauchy numbers at a fixed Strouhal Number *St* = 0.5 are represented on the top-left to bottom-right axis. The bat’s body (absent from flow computation) is represented by the opaque ellipsoid, which is not necessarily scaled to the wing. Arrows identify Leading Edge Vortex (LEV).

The Strouhal number strongly influences the vortex structures generated by the wing (bottom-left to top-right axis in Fig. 3), particularly on the leading edge vortex (LEV) which generates large suction forces on the wing when fully attached [32]. At low flapping speed, a very weak LEV is generated, and the structures shed from the wing’s trailing edge are confined to the tip of the wing, see Fig. 3. Increasing the Strouhal number to *St* = 0.5 generates a stronger LEV indicating much larger force production. A further increase of the Strouhal number to *St* = 0.7 generates a large but unstable LEV which breaks down almost immediately, limiting the duration of effective force production. The tip vortex located on the inner part of the wing results from our simplification of the bat wing, where only the handwing is modeled and is thus not present in real bat flight. However, its strength is many times less than the LEV and tip vortex due to its smaller dynamic velocity (Figure 1d) and it’s influence on the results is minor.

In contrast, variations in the stiffness of the membrane (top-left to bottom-right axis in Fig. 3) has a smaller effect on the bat’s wake. For stiff membranes (i.e. high *Ca*) a LEV is generated on the wing’s leading edge, but the vortices shed at the trailing edge are not spanning the entire wing span. A reduction of the membrane’s stiffness (i.e smaller *Ca*) allows recovering strong vortices spanning the entire wing at the trailing edge, but a very elastic membrane generates a moderately unstable LEV. This moderately reduces the effective force production, similar to (but less extreme than) the high *St* case.

Overall, the wake is governed by the flapping speed of the wing, but within a fixed Strouhal number, the membrane’s stiffness can further influence the vortex structures. We note that our simulation are able to capture some wake features observed experimentally [33], such as the span reduction during the upstroke and the transition vortex (.i.e the vortex generated during the upstroke-downstroke transition), see the electronic supplementary material video.

### Effect of Strouhal number

Further quantification of the results are presented in terms of thrust, lift and power coefficient

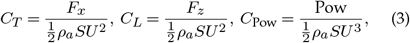

where measured thrust force (*F*_*x*_), lift force (*F*_*z*_), and the power (Pow) are calculated from the integration of pressure and viscous forces over the wing. *S* is the planform area of the wing, and *U* is the forward speed of the bat. To quantify the thrust and lift efficiency of the wing, we use the propulsive and *Rankine-Froude* (RF) [34] efficiency, respectively

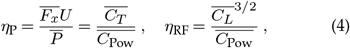

where an overline signifies cycle-averaged values.

The propulsive and lift efficiency for two linear isotropic membranes with different *Ca* for a range of Strouhal numbers are presented in Fig. 4A. Regardless of membrane stiffness, both efficiencies peak at a Strouhal number *St ∼*0.5. The reduced efficiency at high (*St* = 0.7) and low (*St* = 0.3) Strouhal numbers can be associated with the high mixing present in the wake and the small leading edge vortex present on the tip of the wing and not the entire span, see Fig. 3. Peak efficiencies are associated with a strong leading edge vortex, spanning the entirety of the wing span and a small amount of mixing in the wake.

**Figure 4.**
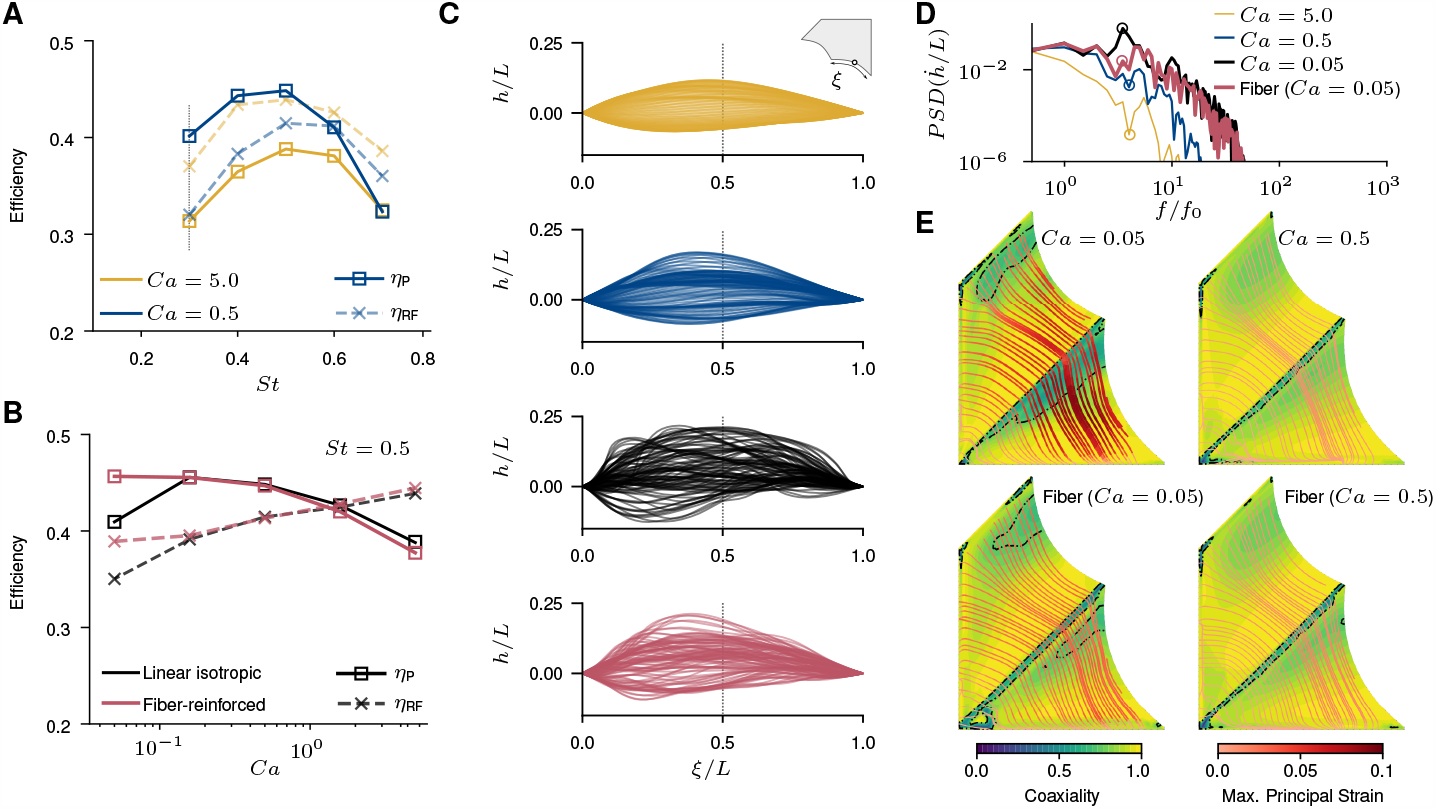
Propuslive and Rankine-Froude efficiency for; **A** a linear isotropic membrane wing at various Strouhal numbers and two Cauchy numbers, and **B** for various Cauchy numbers and membrane material models at a fixed Strouhal number of *St* = 05. The dashed vertical line in A indicates the optimal propulsive efficiency for an identical wing model under symmetric-flapping kinematics. **C** Overlay of the phase-averaged vertical deformation of the trailing edge of the inner membrane at 100 times over a flapping cycle shown in the curvilinear coordinate system of the undeflected membrane trailing edge (see insert). Deflections are shown for three Cauchy numbers for the linear isotropic membrane (*Ca ∈ {*5.0, 0.5, 0.05*}*) and a fiber-reinforced membrane (with an equivalent *Ca* = 0.05). **D** Power-spectral density of the vertical velocity of the midpoint of the membrane’s trailing edge against normalized frequency for the four material properties shown in C. Fiber-reinforced membranes reduced the peak frequency magnitude tenfold compared to equivalent linear elastic membranes. **E** Streamlines of the direction of the max. principal strain ***ε***_1_ colored by the magnitude of the max. principal strain in the membrane at *t/T* = 0.78, corresponding to the flow plot in Fig. 3, for two different constitutive models and membrane Cauchy numbers. Isocontours show the coaxialiality of max. principal strain with the max. principal stress ***σ***_1_. The use of fibers reduces the regions of the membrane where coaxiality *<* 0.75, indicated by the dashed contour. Contours and streamlines are shown in the undeformed configuration.

The efficiency is much less sensitive to membrane stiffness than to *St*. The lift efficiency decreases slightly with the 10x decrease in stiffness, measuring around 10% lower for *Ca* = 0.5 than for *Ca* = 5.0 across the *St* range. In contrast, the thrust efficiency increases with reduced stiffness, with a 25% improvement at *St* = 0.5 (although no change is measured for *St >* 0.6). In terms of thrust coefficient, we find that *Ca* = 0.5 out-performs *Ca* = 5.0 for *St <* 0.6, while the opposite is true for *St >* 0.6. Lift steadily increases with *St* for both membrane stiffness but at a lower rate for *Ca* = 0.5, especially for *St >* 0.6. This is likely linked to a reduction of the effective angle of attack of the wing as *St* increases, which results in less membrane camber and less force generation for *Ca* = 0.5.

The Strouhal number corresponding to the peak in efficiency is relatively insensitive to the membrane stiffness but is strongly influenced by the kinematics. Indeed, replicating these simulations with identical geometrical and material models under *symmetric-flapping* kinematics, which are based on our bat kinematics but use symmetric pitching and flapping of the wing, showed a peak efficiency at *St ∼* 0.3, see SI Appendix Fig. S3. While geometric factors, such as the propulsor’s aspect ratio, have been known to change the optimal Strouhal number [35], our results show that keeping identical material and geometrical models but changing the motion to bat kinematics results in a 66% increase in the optimal flapping rate.

### Effect of membrane stiffness and fiber reinforcement

At the peak-performance Strouhal number (*St* = 0.5), reducing membrane stiffness improves propulsive efficiency while only slightly penalizing the lift efficiency (Fig. 4A). Fig. 4B shows the change in the efficiency at *St* = 0.5 over a 100x reduction in membrane stiffness to quantify the performance of a highly compliant membrane. Additionally, actual bat membranes are made of complex fiber arrangements [17], and the mechanical properties deviate significantly from those of linear isotropic elastic materials, see Fig. 2C. To investigate the effect of the fiber reinforcement on bat flight, we repeat these simulations using a fiber-reinforced membrane, see Materials and Methods; the results are shown in Fig. 4B on top of the linear isotropic elastic membrane results. For linear isotropic membranes, the propulsive efficiency increases with reduced membrane stiffness (i.e. reduced *Ca*). The maximum propulsive efficiency is reached at *Ca ∼* 0.1 before abruptly dropping. Lift efficiency steadily decreases with decreased membrane stiffness, and the drop observed *Ca <* 0.1 for the propulsive efficiency is not as severe. Adding fiber reinforcements to the membranes does not significantly change the performance of stiff membranes (i.e. large *Ca*). However, at very low stiffness (i.e. small *Ca*), adding fiber reinforcements allow a 10% gain in propulsive and lift efficiencies and no drop-off. This loss in efficiencies near *Ca* = 0.05 for linear isotropic membranes corresponds to the onset of flutter of the membrane and increased vortex shedding in the wake, see Fig. 3. Figure 4C documents the flutter in terms of the phase-averaged trailing edge deflection of the membrane. Deflections are shown in the frame of reference of the undeformed membrane (see insert in Fig. 4C). The corresponding power-spectral density (*PSD*) of the vertical velocity of the trailing edge’s midpoint also indicates flutter, see Fig. 4D. Membrane flutter introduces high-frequency oscillations of the membrane and shifts the peak frequency of the membrane above the motion frequency. This high-frequency oscillation contains ten times the energy of stiffer membranes. By introducing fiber reinforcement to the membranes, the internal response of the wing is changed (see below), and the aerodynamic efficiency is improved.

These very flexible membranes also increase the high-frequency mixing of the fluid, as measured by the *enstrophy ε*, the integral of the square-vorticity in the fluid domain. We measure a strong inverse Pearson’s correlation *ρ*_*ε,η*_ = *−*0.976 with *p*-value of 0.00436 between the mean enstrophy during a cycle and the propulsive efficiency at *St* = 0.5, see SI Appendix Fig. S4. This is because excessive fluid mixing requires significant power input from the wing without a large gain in force production. The flutter induced by a very flexible membrane generates the most enstrophy in the bat’s wake, explaining the poor aerodynamic efficiency in Fig 4B at very low *Ca*, whereas fiber-reinforced membrane limits the flutter and thus the mixing in the wake, which allows maintaining high aerodynamic efficiencies.

We document the internal response of the membrane in terms of the direction and magnitude of the principal strain in Fig. 4E. When the membrane has a low linear stiffness (*Ca* = 0.05), fiber reinforcements greatly reduce the maximum strain in the membrane from 9.4% to 4.1% due to the loading-induced non-linear stiffening of the fiber at high strain, see Fig. 2C. This effect is similar to an increase in stiffness of a linear isotropic membrane without requiring a thicker membrane, which explains the extremely low thickness of the bat’s membrane. This loading-induced stiffening does not activate for the *Ca* = 0.5 membrane because the strains are too low. Fig. 4E also shows that fiber-reinforcing the *Ca* = 0.05 membrane also reduces the region of low coaxiality of the principal stress and strain in the membrane by 60%. Coaxiality of the stress and strain tensors corresponds to a state of deformation in which the principal directions of both tensors are aligned [36]. This metric indicates the state of isotropy of stress and strain that biological tissues appear to maximize through internal microstructure evolution since it minimizes their strain energy [37, 38]. This principle of energy minimization is commonly considered a fundamental axiom in Nature.

## Conclusion

In this paper, we performed fully-coupled computational fluid-structure interaction simulations of a bat wing with fully adjustable parametric kinematics and material model. The parametric models are simplified compared to real-life bat kinematics and wings, but capture the essential nonlinear aspects of bat flight and mechanical membrane behavior (see Fig. 2C-D), making these results relevant to actual bat flight. These models allow us to separately investigate the effects of the Strouhal and membrane compliance on the flight performance of bats, as well as giving detailed insights into the structural response of the wing’s membrane. First, we show that bats operate at a Strouhal number corresponding to a peak in both propulsive and lift efficiencies. This peak occurs near Strouhal number *St ∼* 0.5, which agrees well with the mean Strouhal number of actual bats; see Fig. 1A. This peak is also well above the range commonly associated with optimal locomotion in birds and fish [16]. We demonstrate that this optimum results from specific structures present in the wake of the bat and that these occur at high flapping speeds due to the highly three-dimensional nature of bat kinematics. Indeed, when replicating the simulations under symmetric-flapping kinematics, we find that the optimum shift to the classical *St ∼* 0.3. This implies that the high Strouhal number of our model results from the specialized kinematics, not the material flexibility.

Finally, we show that reducing membrane stiffness benefits propulsive efficiency. However, for very compliant (i.e small *Ca*) linear isotropic membranes, we observe membrane flutter and a severe deterioration of the efficiencies. As a result, a strong inverse correlation (*ρ*_*ε,η*_ = *−*0.976) between mean flow enstrophy (mixing) and propulsive efficiency is found. This suggests that membrane wings made of linear isotropic elastic materials are most efficient just before the onset of flutter, which we estimate to occur around *Ca ∼* 0.1. By reinforcing the isotropic membranes with fibers, which capture the response of actual bat skin more accurately, flutter is delayed and high aerodynamic efficiencies can be maintained for *Ca <* 0.1 . This effect is similar to an increase in membrane stiffness but without the resulting increase in skin thickness and wing mass, which is likely to have a positive influence on the cost of flight [11], and the nonlinear effect is more pronounced when the loading on the membrane is large, i.e. for high-speed flight. This suggests that the complex fiber arrangement in the bat wing’s membrane has evolved to optimize the structural response of the wing by limiting flutter even when flying quickly, ultimately improving its propulsive performance.

## Methods

### Bat flight kinematics

A parametric model for the wing’s motion lets us freely prescribe the bat Strouhal number. Five degree-of-freedom motion is imposed at the carpus and the rest of the wing passively deforms under aerodynamic and inertial loading. Our parametric kinematic model closely matches experimental measurements, see Fig. 2D, capturing key features of bat flapping such as the angled downstroke plane and the biased power and recovery strokes.

The parameterized flight kinematics is expressed in a fixed Cartesian or global coordinate system (*OXYZ*) and describes the motion and orientation of the carpus (*oxyz*) coordinate system, see Fig. 2B.

The time-dependent angles and translation applied to the carpus are given by

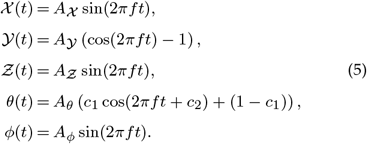

where 𝒳(*t*) is the surge, *𝒴*(*t*) the sway, 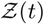 the heave, *θ*(*t*) the pitch and *ϕ*(*t*) the roll. Pitch and roll are rotations around the *Y* and *X*-axis, respectively. The non-dimensional amplitudes are *A*_𝒳_/*b* = *A*_𝒴_/*b* = 0.10 and *A*_*𝒳*_ */b* = 0.15 where *b* is the wing span. These motion amplitudes have been derived from a fit of motion of the wing such that the stroke plane angle is *∼* 60^*°*^ from the horizontal, typical of medium speed flight of bats [4] for a *Glossophaga soricina*, or Palla’s long-tongued bat. The roll amplitude is *A*_*ϕ*_ = 30^*°*^, taken from the mean stroke amplitude in [13]. Other authors have reported similar kinematics [4], see Fig. 2. The contraction/extension of the wing span during the down/up-stroke, resulting from the variation of the *𝒴* position of the carpus is *A*_*𝒴*_*/b ∼* 10%.

### Optimal Pitch Profile

The effective angle of attack measures the angle between the flow and the wing at the 3*/*4 chord, *α*_3*/*4_, during motion and includes a dynamic upwash correction. For more details, the reader is referred to [28]. To model the different flow conditions along the span, we apply a strip-theory approach, where the wing is discretized in 10 spanwise strips of equal width. For each strip, we compute the effective angle of attack of a point located at 3*/*4 chord in the middle of the span of the strip. The total effective angle of attack of the wing is found by an area-weighed average of the different strips.

Although this method cannot accurately predict the forces during a cycle, it allows us to estimate the effective angle of attack seen by the wing during a cycle. We can now optimize our analytic form for *θ*(*t*) given by Equ. 5 to achieve the target angle of attack profile 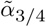 by minimizing a constrained equation of the form

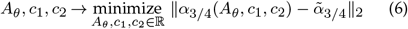

with *A*_*θ*_ *∈* [0, *π/*2], *c*_1_ *∈* [0.5, 1] and *c*_2_ *∈* [*−π/*2, *π/*2]. We use a simple form for the target effective angle of attack, where the upstroke effective angle of attack must vanish and reach target value *α*_*max*_ during the downstroke

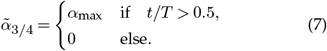

We set *α*_max_ = 20; a common mean effective angle of attack during downstroke [39]. The minimization procedure is carried out for each Strouhal number and results in various amplitudes *A*_*θ*_, but the other two values were constantly around *c*_1_ *∼* 0.8 and *c*_2_ *∼ −π/*8. Therefore, we fixed those values and repeated the optimization for *A*_*θ*_ only, resulting in *A*_*θ*_ = [0.28, 0.45, 0.63, 0.80, 0.98] radians for *St* = [0.3, 0.4, 0.5, 0.6, 0.7], respectively. Sample results of the optimization are presented in SI Appendix Fig.S2, showing that this method generates roughly constant lift during the down-stroke and very small lift in the upstroke, consistent with actual bat flight [9].

### Hyperelastic formulation

To investigate the effect of membrane compliance and fiber reinforcement on flight efficiency, we use two constitutive models for the membrane; a linear isotropic elastic model and a hyperelastic fiber-reinforced model calibrated on experimentally obtained bat wing membrane properties [18, 31], see Fig. 2C. The fiber-reinforced model introduces microstructural and loading-induced anisotropy in the response of the membrane via a deformation invariant-based hyperelastic strain energy density function [40]. This type of constitutive approach is suited to model a wide class of fibrous biological soft tissues [41–43], including bat wing membrane [31].

Bones in the handwing are more flexible than the humerus and radius and are orders of magnitude stiffer than the membrane they support. In this work, we model the digits with an isotropic material whose Cauchy number is 3000 times larger than the stiffest membrane (i.e. *Ca* = 5), making it it effectively highly rigid compared to the membrane. Additionally, the Cauchy scaling using *U*_ref_ ensures that the bone’s deformation is constant (and minimal) across Strouhal numbers. The camber in the find is thus only generated by the deformation of the wing’s membrane. Digits typically have thicknesses much greater than the membrane they support. Our simplified model uses a uniform thickness throughout the whole wing and scales the bone’s stiffness and density accordingly. To model the fiber-reinforced membrane of the wing, we use a constitutive model based on the strain energy density function proposed by [40, 44]

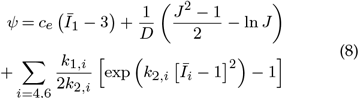

where *c*_*e*_ is a material constant related to the matrix shear modulus *μ* by *c*_*e*_ = *μ/*2 and *D* is a material parameter related to the matrix compressibility, or the bulk modulus *K* = 2*/D. I*_1_ is the first invariant of the isochoric right Cauchy-Green deformation tensor 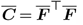, with 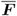 the isochoric part of the deformation gradient *F* calculated as 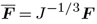, where *J* is the Jacobian or determinant of the deformation gradient *F*. The anisotropic response of the fiber is introduced through the undeformed mean fiber vector a_*i*_ and the pseudo-invariant of 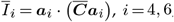, and represents the stretch along each fiber direction (in this case we have two fiber families, denoted by their invariant-associated index, 4 and 6). *k*_2*i*_ is a dimensionless positive parameter controlling the shape of the nonlinear stiffening of the fibers’ mechanical response, and *k*_1,*i*_ is an elastic modulus-like parameter corresponding to fiber stiffness. The constitutive parameters of this model are obtained by numerical identification from experimentally measured bat wing membrane properties [18].

The membrane material was assumed to be relatively incompressible by setting the compressibility parameter *D* = 20*c*_*e*_, leading to an equivalent ground-state Poisson’s ratio *ν* = 0.475. The 20 ratio of bulk to shear properties was sufficiently small to prevent numerical ill-conditioning associated with volumetric locking in purely displacement-based finite element formulations [45] whilst also ensuring low compressibility.

#### Hyperelastic calibration

To identify the constitutive parameters in Equ. 8, we fit the experimental data from [18, 31] to the stress-strain curves from the model in 8 under the same equibiaxial loading.

The coefficients [*c*_*e*_, *k*_1,4_, *k*_2,4_, *k*_1,6_, *k*_2,6_] are determined by a *L*_2_-norm minimization of the difference in the stress-strain curves *σ*_*ii*_ and the experimental stress/strain curve 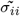

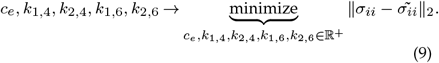

In practice, we find that *c*_*e*_ tends to take negative values, as this would result in unphysical material behavior, we fix its value to *c*_*e*_ = 20. The different parameters found are *k*_1,4_ = 33.5, *k*_2,4_ = 13.5, *k*_1,6_ = 102.7, *k*_2,6_ = 12.3.

These coefficients can then be injected into the numerical model for the bat wing; the difficulty lies in scaling the fitted parameters into a dimensionless form employed in our numerical simulation. To scale the fitted hyperelastic parameters and compare fiber-reinforced membranes with isotropic membranes, we must be able to express both in terms of *Ca*. For a linear isotropic elastic constitutive law, *E* is readily available (and constant). For the transversely isotropic hyperelastic fiber-reinforced membrane, *E* is a function of the deformation (through 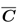), making a simple estimate unavailable. To alleviate this problem, we use an initial tangent stiffness modulus approach to estimate the initial stiffness of the hyperelastic material (in the limit of vanishing strain), see Fig. 2C. This tangent stiffness modulus approach ensures that linear isotropic and non-linear fiber-reinforced membranes behave similarly in the small strain limit and ultimately allow us to compare them. Finally, we use Equ. 2 to express the different material coefficients at different Cauchy numbers.

### Numerical Setup

We perform all the simulations with our validated fluid-structure interaction solver [29, 30] which couples an immersed boundary finite volume method for the fluid with a shell finite element method for the structure. Defining *b* as the hand-wing span, the fluid domain consists of a uniform region of dimensions [2.5, 1.2, 1.2] *× b*, centered around the wing. Grid stretching fills the domain until it reaches a total size of [12, 6, 6] *× b*. Because of the symmetry of the problem, we only model the bat’s right wing and apply a symmetry boundary condition on the centerline of the domain (i.e., the middle of the bat’s body) to reduce the computational load. A uniform free-stream velocity *U* is imposed on the inlet of the fluid domain, while a zero pressure gradient condition is used on the outlet. The no-slip condition is applied to the immersed wing, and all other fluid domain boundaries are threaded as free-slip walls. The wing is modeled as an initially flat shell with constant thickness *h/b* = 0.005 and the fixed-point fluid-structure interaction problem is solved using a quasi-Newton scheme until the relative residuals between consecutive coupling iterations drop below 10^*−*4^ for both the displacements and the forces [30]. This typically takes 2 iterations per time step, as the mass ratio of the simulations is relatively large (small added-mass effects). In our simulations, the flow evolves for six cycles. However, we found excellent repeatability after two cycles and we phase-average the quantities of interest over the last four cycles. The cycle-averaged values are then obtained by averaging the phase-averaged data.

Table 1 shows the results of a mesh convergence study for a Strouhal number *St* = 0.5 and a Cauchy number *Ca* = 0.5. We vary the mesh resolution *N*_*x*_ = *b/Δx ∈* [32, 64, 96, 128]. We compare the cycle-averaged forces coefficients to the finest mesh and find that reducing the resolution toward *N*_*x*_ = 128 changes the resulting forces by *∼* 1%. We use the *N*_*x*_ = 96 mesh for all the simulations presented herein, giving a non-dimensional wall distance *y*^+^ *∼* 5, assuming a flat plate 1/7 power-law for the velocity profile. The total fluid mesh count is around 12.8M control volumes. For the structural model, 64 linear triangular elements along each digit, giving a total mesh count of 13,093 elements.

**Table 1.**
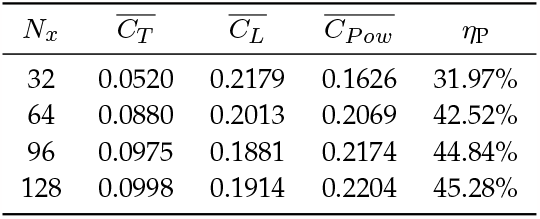
Convergence of aerodynamic force coefficients of the bat wing simulations for a Strouhal number *St* = 0.5 and *Ca* = 0.5 using a isotropic linear elastic membrane for various resolutions.

| $N_x$ | $\overline{C_T}$ | $\overline{C_L}$ | $\overline{C_{Pow}}$ | $\eta_P$ |
| --- | --- | --- | --- | --- |
| 32 | 0.0520 | 0.2179 | 0.1626 | 31.97% |
| 64 | 0.0880 | 0.2013 | 0.2069 | 42.52% |
| 96 | 0.0975 | 0.1881 | 0.2174 | 44.84% |
| 128 | 0.0998 | 0.1914 | 0.2204 | 45.28% |

Simulations were performed on the *Iridis 5* supercomputer at the University of Southampton. Typical simulations used 64 2.0-GHz Intel Skylake processors for the fluid domain and 4 for the structural problem. Simulations took 3-4 days to reach six motion cycles (at *St* = 0.5) at this resolution with a fixed time step *ΔtU/Δx* = 0.2.

## Data Accessibility

The data generated during this study and the analysis scripts have been uploaded to 10.4121/b6bb4c63-dc61-4a7d-9ad9-66f82139604a.v1.

## Authors’ Contributions

M.L., G.D.W., and G.L. designed the research; M.L. performed the research; M.L., G.D.W. and G.L. analyzed the data; and M.L, G.D.W. and G.L. wrote the paper. All authors gave final approval for publication and agreed to be held accountable for the work performed therein.

## Acknowledgements

We would like to thank Kenny S. Breuer for comments on an early version of this manuscript.

