## Supplementary material for "Rapid flapping and fiber-reinforced membrane wings are key to high-performance bat flight": JRSI_SI_accepted.pdf

This document contains the Supplementary Information for the article "Rapid flapping and fiber-reinforced membrane wings are key to high-performance bat flight".

#### Wake width length scale

In figure S1, we compare the wake width (.i.e. vertical, or $z$-direction, dimension of the wake) of our bat model flying at different Strouhal numbers. The wake width is obtained by displaying isosurfaces of $\lambda_2$-criterion at the peak motion amplitude ($A$). We find that the tip-to-tip amplitude is an excellent measure of the wake width generated by the bat. This justifies its use in the definition of the Strouhal number.

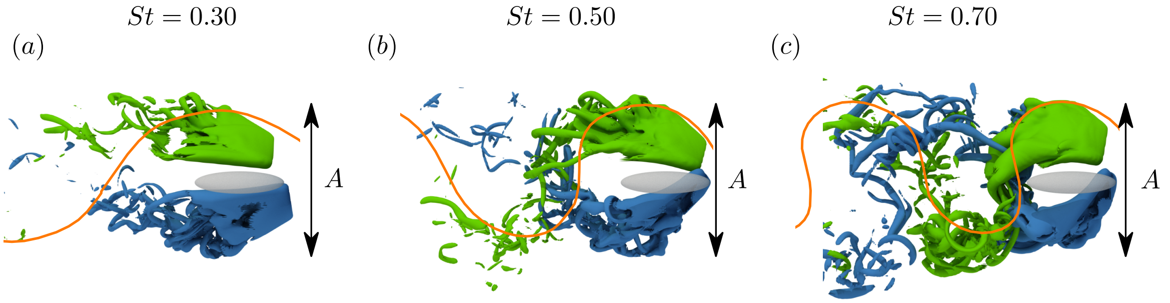


Figure S1: Overlaid wake, represented as isosurfaces of $\lambda_2$-criterion ($\lambda_2(L/U)^2 = -1.2 \times 10^{-5}$), from two phases of the cycle (start and end of the downstroke are colored green and blue, respectively) to illustrate the maximum extensions of the wake for three Strouhal numbers. The grey-shaded ellipsoid represents the bat's body. The bat is viewed from the right side, flying forward from left to right. The right wing has been omitted for clarity. The orange line depicts the wing-tip trace during flapping.

### Experimental data of bat flight

In Table S1, we present the detailed data from [1] that we rearranged into the standard Strouhal

$$St = \frac{fA}{U}, \tag{1}$$

where $f$ id the flapping frequency, $U$ is the constant forward speed of the bat and $A$ is the peak-to-peak motion amplitude, see previous section. The different symbols correspond to: $N$, the number of individual animals tested for each species; $n$, the number of test points for each species; $t$, the total number of wingbeat cycles recorded with amplitude data; $c$, the mean number of wingbeat cycles per test point. Values for $f_w$ and $\theta_w$ are means ±S.D. with the S.D. computed as $\frac{1}{N}\sqrt{\sum^N \sigma^2}$. The Strouhal range has been calculated using the mean wingbeat amplitude and frequency and the speed range to obtain a min/max Strouhal range.

Table S1: Summary of flight characteristics for different species of bats, adapted from [1]. The different symbols correspond to: $N$, the number of individual animals tested for each species; $n$, the number of test points for each species; $t$, the total number of wingbeat cycles recorded with amplitude data; $c$, the mean number of wingbeat cycles per test point. Values for $f_w$ and $\theta_w$ are means ±S.D. with the S.D. computed as $\frac{1}{N}\sqrt{\sum^N \sigma^2}$. The Strouhal range has been calculated using the mean wingbeat amplitude and frequency and the speed range to obtain a min/max Strouhal range.

| Species | $N$ | $n$ | $t$ | $c$ | Wing span (m) | Speed range ($ms^{-1}$) | Wingbeat frequency $f_w$ (Hz) | Wingbeat amplitude $\theta_w$ (degrees) | Strouhal range (-) |
|---|---|---|---|---|---|---|---|---|---|
| *Chalinolobus gouldii* | 40 | 70 | 244 | 6.10 | 0.3457 | 3.3-10.0 | 9.04±0.87 | 65.48±31.56 | 0.54-0.18 |
| *Chalinolobus morio* | 4 | 14 | 118 | 6.94 | 0.2877 | 2.2-7.8 | 10.91±1.01 | 49.23±15.39 | 0.61-0.17 |
| *Chalinolobus nigrogriseus* | 4 | 4 | 35 | 8.75 | 0.278 | 5.6-9.4 | 11.27±0.84 | 56.67±15.28 | 0.28-0.16 |
| *Hipposideros ater* | 9 | 20 | 75 | 8.33 | 0.2489 | 2.2-4.7 | 10.91±0.8 | 57.14±8.88 | 0.62-0.29 |
| *Macroderma gigas* | 1 | 8 | 60 | 6.67 | 0.759 | 2.8-8.1 | 6.69±0.71 | 83.75±18.47 | 1.33-0.46 |
| *Miniopterus schreibersii* | 1 | 2 | 25 | 12.50 | 0.3409 | 5.8-8.9 | 9.1±0.57 | 92.5±17.68 | 0.43-0.28 |
| *Mormopterus planiceps* | 34 | 72 | 211 | 6.81 | 0.2635 | 2.1-9.4 | 9.34±1.38 | 40.68±10.15 | 0.42-0.09 |
| *Nyctophilus arnhemensis* | 2 | 31 | 259 | 8.35 | 0.3014 | 1.1-4.7 | 11.43±1.13 | 33.44±11.18 | 0.91-0.21 |
| *Nyctophilus geoffroyi* | 6 | 23 | 163 | 9.88 | 0.2631 | 0.8-7.2 | 10.94±0.96 | 43.92±16.44 | 1.38-0.15 |
| *Nyctophilus gouldi* | 6 | 22 | 221 | 10.52 | 0.3046 | 1.7-5.8 | 10.4±1.05 | 59.29±10.99 | 0.96-0.28 |
| *Nyctophilus timoriensis* | 4 | 17 | 224 | 13.93 | 0.3219 | 1.4-6.7 | 10.56±0.59 | 48.8±18.74 | 1.03-0.22 |
| *Nyctophilus timoriensis* | 2 | 10 | 150 | 15.00 | 0.3495 | 1.7-2.5 | 11.08±0.34 | 47.5±5.4 | 0.94-0.64 |
| *Pteropus poliocephalus* | 3 | 15 | 21 | 7.00 | 1.338 | 3.1-8.6 | 3.4±0.88 | 86.67±12.58 | 1.11-0.40 |
| *Pteropus scapulatus* | 7 | 7 | 40 | 5.71 | 1.106 | 6.7-10.0 | 4.15±0.5 | 94.29±17.18 | 0.56-0.38 |
| *Rhinonycteris aurantius* | 5 | 13 | 156 | 9.18 | 0.308 | 2.5-7.2 | 9.76±0.55 | 70.77±14.12 | 0.74-0.26 |
| *Saccolaimus flaviventris* | 1 | 12 | 91 | 5.92 | 0.575 | 1.7-5.3 | 8.36±0.75 | 48.33±8.66 | 1.19-0.38 |
| *Scotorepens balstoni* | 1 | 9 | 64 | 10.69 | 0.266 | 3.3-5.0 | 11.31±0.67 | 35±8.29 | 0.28-0.18 |
| *Scotorepens greyi* | 7 | 7 | 80 | 13.00 | 0.25 | 6.7-8.3 | 11.59±1.01 | 60±18.26 | 0.23-0.18 |
| *Tadarida australis* | 14 | 23 | 154 | 9.60 | 0.4625 | 3.6-13.5 | 8.19±1.08 | 90.74±27.7 | 0.83-0.22 |
| *Taphozous georgianus* | 1 | 5 | 44 | 8.80 | 0.4637 | 2.5-4.2 | 8±1.89 | 71±7.42 | 0.92-0.55 |
| *Taphozous hilli* | 6 | 6 | 58 | 8.29 | 0.4616 | 6.1-8.3 | 7.47±0.46 | 100±8.94 | 0.49-0.36 |
| *Vespadelus finlaysoni* | 12 | 12 | 88 | 6.77 | 0.2549 | 4.7-8.1 | 10.68±1.11 | 58.18±9.82 | 0.29-0.17 |
| *Vespadelus regulus* | 7 | 14 | 145 | 10.52 | 0.2335 | 1.4-6.9 | 10.75±0.57 | 41.38±6.85 | 0.65-0.13 |
| Average | | | | | 0.349 | 3.010-7.238 | 9.89±0.87 | 59.71±13.82 | 0.72-0.27 |

### Optimal pitch profile

This section presents the result of adjusting the pitch profile on the aerodynamic performances.

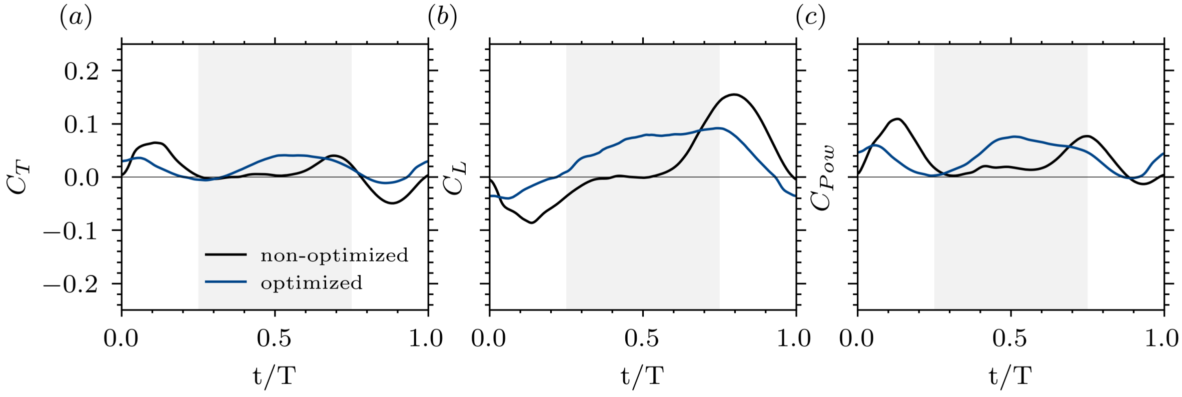


Figure S2: Phase-averaged (*a*) Thrust, (*b*) Lift and (*c*) Power coefficient for two wings at $St = 0.5$ under different pitch profiles. A non-optimized pitch profile (using $A_\theta, c_1, c_2 = 0.8, 1, 0$) generates inconsistent forces for during the downstroke (represented as a gray-shaded area), and the peak power consumption occurs in the upstroke when the bat should be "feathering". The optimized pitch profile results in a power stroke, where the forces are predominantly produced in the downstroke and the power during upstroke is minimized.

### Symmetric-flapping kinematics and results

We compare our bat kinematics to symmetric flapping kinematics, where the wing only rolls and pitches around the shoulder. We investigate simple symmetric propulsive flapping without attempting to generate efficient lift, given by

$$\begin{aligned}\mathcal{X}(t) &= A_{\mathcal{X}} \sin(2\pi f t),\\ \mathcal{Y}(t) &= A_{\mathcal{Y}} \sin^2(2\pi f t),\\ \mathcal{Z}(t) &= A_{\mathcal{Z}} \sin(2\pi f t),\\ \theta(t) &= A_{\theta} \cos(2\pi f t),\\ \phi(t) &= A_{\phi} \sin(2\pi f t).\end{aligned} \tag{2}$$

where $\mathcal{X}(t)$ is the surge, $\mathcal{Y}(t)$ the sway, $\mathcal{Z}(t)$ the heave, $\theta(t)$ the pitch and $\phi(t)$ the roll. Pitch and roll are rotations around the $Y$ and $X$-axis, respectively. The non-dimensional amplitudes are $A_{\mathcal{X}}/L = 0.10$, $A_{\mathcal{Y}}/L = -0.15$ and $A_{\mathcal{Z}}/L = 0.15$ where $L$ is the wing span. The pitch and roll amplitude are taken as typical values[3], $A_{\theta}/L = 20$ and $A_{\phi}/L = 30$, respectively. These kinematics were applied to the same bat handwing model, using the linear isotropic membrane with $Ca = 0.5$, and the resulting thrust, lift, power coefficients, and the resulting propulsive efficiency are presented in Fig. S3. In contrast to the propulsive peak of $St = 0.5$ for the bat kinematics, flapping the flexible bat wing with these symmetric-flapping kinematics results in a propulsive efficiency peak at $St = 0.3$, agreeing with the theorized peak efficiency range of $0.25 - 0.35$[2].

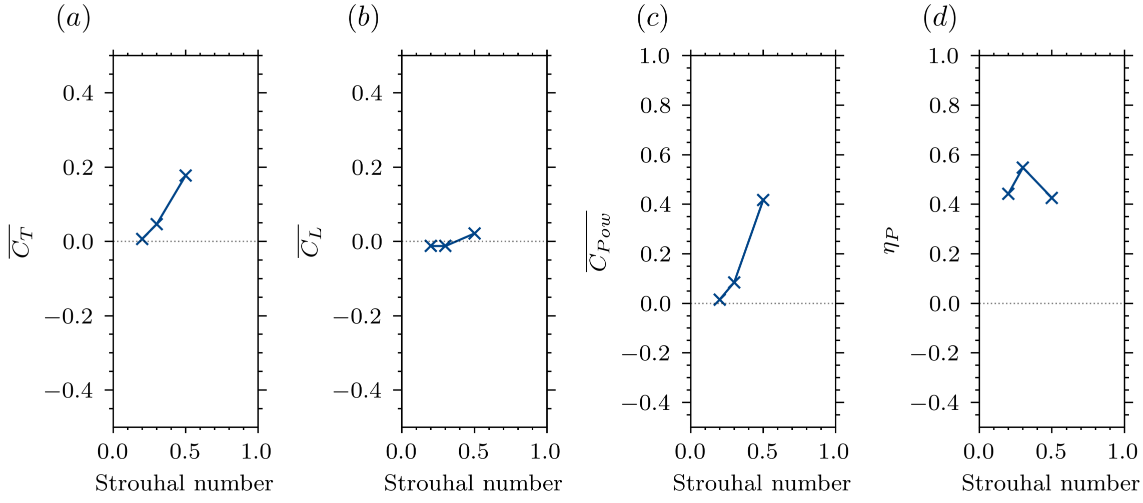


Figure S3: Cycle-average thrust ($a$), lift ($b$) and power coefficient ($c$) for a linear isotropic elastic wing at a Cauchy number $Ca = 0.5$ under bird-like kinematics for a range of Strouhal number. Resulting propulsive efficiency ($d$), peaking at $St = 0.3$.

### Enstrophy - efficiency correlation

We compute the (instantaneous) fluid enstrophy in the fluid domain as the integral of the square vorticity

$$\mathcal{E} = \frac{1}{2}\int_{\text{Fluid}} |\boldsymbol{\omega}|^2 \, \mathrm{d}\boldsymbol{x}_f \tag{3}$$

were $\boldsymbol{\omega}$ is the vorticity vector and $\boldsymbol{x}_f$ is a point in the fluid domain. Next, the *Pearson correlation coefficient* between the mean of the phase-averaged total enstrophy ($\bar{\mathcal{E}}$) in the domain and the resulting propulsive efficiency is given by

$$\rho_{\bar{\mathcal{E}},\eta} = \frac{\text{cov}(\bar{\mathcal{E}}, \eta_{\text{P}})}{\sigma_{\bar{\mathcal{E}}\sigma_{\eta_{\text{P}}}}} = -0.976. \tag{4}$$

where $\sigma_a$ is the standard deviation of variable $a$, and cov(..., ...) is the covariance. The correlation is associated with a $p$-value of $p = 0.00436$ (significant). This is summarized in Fig. S4.

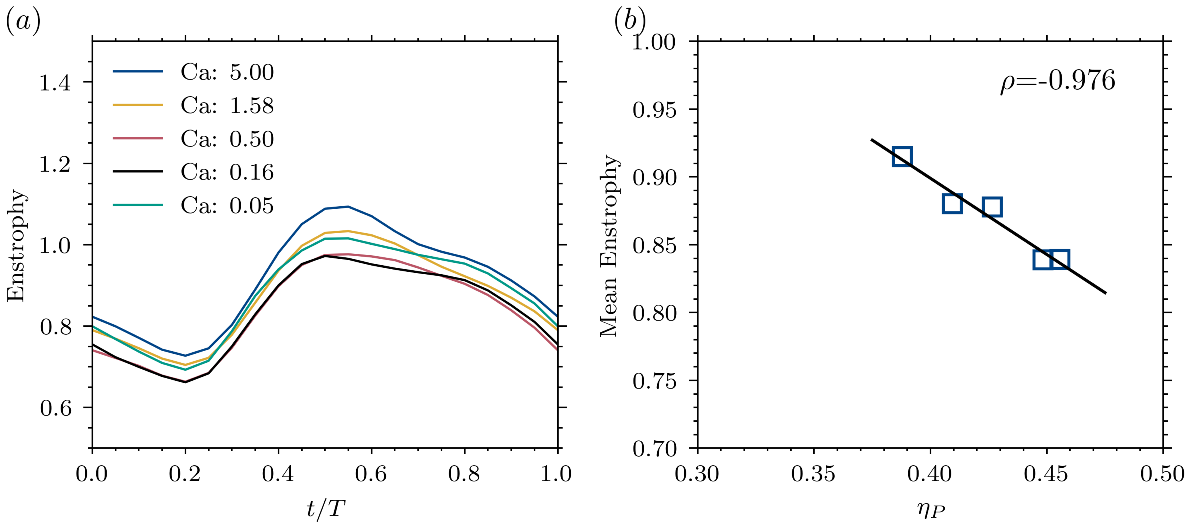


Figure S4: (*a*) Phase-averaged enstrophy for a linear isotropic elastic membrane wing at different membrane Cauchy numbers. (*b*) Pearson correlation coefficient and line of best-fit between propulsive efficiency and mean enstrophy during a cycle for various membrane Cauchy numbers.

#### Principal stress-strain and coaxilaity

For a given stress tensor $\boldsymbol{\sigma}$ (resp. strain tensor $\boldsymbol{\varepsilon}$), there exists a plane with normal $\hat{\boldsymbol{n}}$, such that the shear stresses (resp. shear strain) vanishes. This plane defines one of the principal directions associated with a principal stress (resp principal strain) of the stress (resp strain) tensor. This can be expressed as

$$\boldsymbol{\sigma}\hat{\boldsymbol{n}} = \lambda\hat{\boldsymbol{n}} \tag{5}$$

which defines the following eigenvalue problem

$$(\boldsymbol{\sigma} - \lambda\boldsymbol{I})\,\hat{\boldsymbol{n}} = 0 \tag{6}$$

for which a solution exists only if $\hat{\boldsymbol{n}} \neq 0$ and corresponds to

$$\det(\boldsymbol{\sigma} - \lambda\boldsymbol{I}) = 0. \tag{7}$$

The solution to this eigenvalue problem gives three eigenvalues $\lambda_1 > \lambda_2 > \lambda_3$ and the corresponding three eigenvector $\hat{\boldsymbol{\sigma}}_1$, $\hat{\boldsymbol{\sigma}}_2$, $\hat{\boldsymbol{\sigma}}_2$. These eigenvalues/eigenvectors correspond to the magnitude and direction of the maximum, mean and minimum normal stresses (resp strains).

The coaxiality of the maximum principal stress and maximum principal strain is given by

$$C \equiv \hat{\boldsymbol{\sigma}}_1 \cdot \hat{\boldsymbol{\varepsilon}}_1, \tag{8}$$

A coaxially of 1 implies perfect alignment of the principal stress and principal strains; zero implies that the principal stress and strain are orthogonal.
